# Selection of a Malignant Subpopulation from a Colorectal Cancer Cell Line

**DOI:** 10.1101/578823

**Authors:** Pei-Lun Lai, Ting-Chun Chen, Chun-Yen Feng, Hsuan Lin, Ng Wu, Yun Chen, Michael Hsiao, Jean Lu, Hsiao-Chun Huang

## Abstract

Colorectal cancer (CRC) is a leading cause of death from cancer worldwide. Thus, there is an emerging need for new experimental models that allow identification and validation of biomarkers for CRC-specific progression. In this study, we propose a repeated sphere-forming assay as a strategy to select a malignant subpopulation from a CRC line, HCT116. We validated our assay by confirming that three canonical stemness markers, Nanog, Oct4, and Lgr5, were up-regulated in the sphere state at every generation of the selection assay. The resulting line, after eight rounds of selection, exhibited an increased sphere-forming capacity *in vitro* and tumorgenicity *in vivo*. Furthermore, dipeptidase 1 (DPEP1) was identified as the major differentially expressed gene in the selected clone, and depletion of DPEP1 suppressed the elevated sphere-forming capacity *in vitro* and tumorgenicity *in vivo*. Overall, we have established an experimental strategy for the isolation of a malignant subpopulation from a CRC cell line. Results from our model also suggested that DPEP1 can serve as a promising prognostic biomarker for CRC.

## Introduction

Colorectal cancer (CRC) is the fourth most common cause of cancer death worldwide and is also a common neoplastic disease in modern countries^1, 2^. As with many other types of cancers, the development of CRC from benign adenomas to malignant carcinomas is thought to result from long-term accumulation of mutations during the course of disease progression^3^. Because the survival of CRC patients is closely linked to the time of diagnosis and stage of the tumor^4^, it is increasingly important to identify specific factors involved in CRC progression to serve as prognostic markers. A good experimental model that spans a wide range of malignancies would facilitate identification and validation of such CRC markers.

It has been proposed that cancer stem cells (CSCs), a subpopulation of cancer cells that has self-renewing features to promote tumor growth and resistance to chemotherapy, are the most malignant subset of cells responsible for the recurrence and metastasis of CRC^5, 6^. The sphere-forming assay is the gold-standard for isolating CSC-like cells. The history of the assay can be traced back to the late 1960s when it was used to study neurogenesis in neural stem cells^7^. Specifically, the sphere-forming assay was used to identify cells with higher neurogenic potentials both in clonality and multipotent differentiation^8^. Since then, this assay has been employed to investigate stem cells in a variety of normal tissues^8^. Scientists have adopted the sphere-forming assay to form tumorspheres in many kinds of cancers, including brain^9^, breast^10^, and colorectal^11, 12^. These tumorspheres reportedly have similar self-renewal characteristics and express the same canonical stemness markers (such as Nanog^13^) as normal stem cells. A caveat, however, exists for using tumorspheres, or CSC-like cells, as platforms to study CRC progression. Because tumorspheres are present acutely in a different physiological state (apart from the long-term adherent culture), the measured phenotypes may only reflect transient^14^, and not, stable properties of the cells.

Inspired by such limitations, we used a repeated sphere-forming assay as a strategy to select a malignant cell line that was phenotypically stable. This is conceptually parallel to our previous establishment of a series of metastatic cell lines using repeated invasion assays^15, 16^. Such model cell lines have been used to identify genes or cellular phenotypes that are associated with metastasis^16, 17^. In this study, with a repeated sphere-forming assay, we aimed to select a malignant clone from a HCT116 CRC cell line. Using RNA-seq to compare the transcriptome between the selected clone and its parental cell line, we also identified the gene responsible for malignancy both *in vitro* and *in vivo*.

## Results

### Evolution and selection of a cancer cell line using repeated sphere-forming assays

Our experimental design is illustrated in Figure 1. With an ultra-low attachment dish, we generated tumorspheres using the sphere-forming assay. While the majority of cells died during culturing, a small population (i.e., CSC-like cells) was able to form spheres. These sphere-derived CSC-like cells (SDCSCs) were isolated, and a portion was collected and subjected to experimental validations. The remaining cells were dissociated and re-plated onto a regular dish as adherent cells (sphere-derived adherent cells, or SDACs). SDACs were allowed to expand, some were frozen in stock vials, and after a recovery of 14 days, cells were again cultured in an ultra-low attachment dish to form spheres for the next round of selection (Figure 1). For simplicity, we abbreviated ‘generation’ as ‘G’, ‘SDCSCs’ as ‘S’, and ‘SDACs’ as ‘D’. For example, first-generation SDCSCs are designated as ‘G1S’, and first-generation SDACs are designated as ‘G1D’.

**Figure 1.**
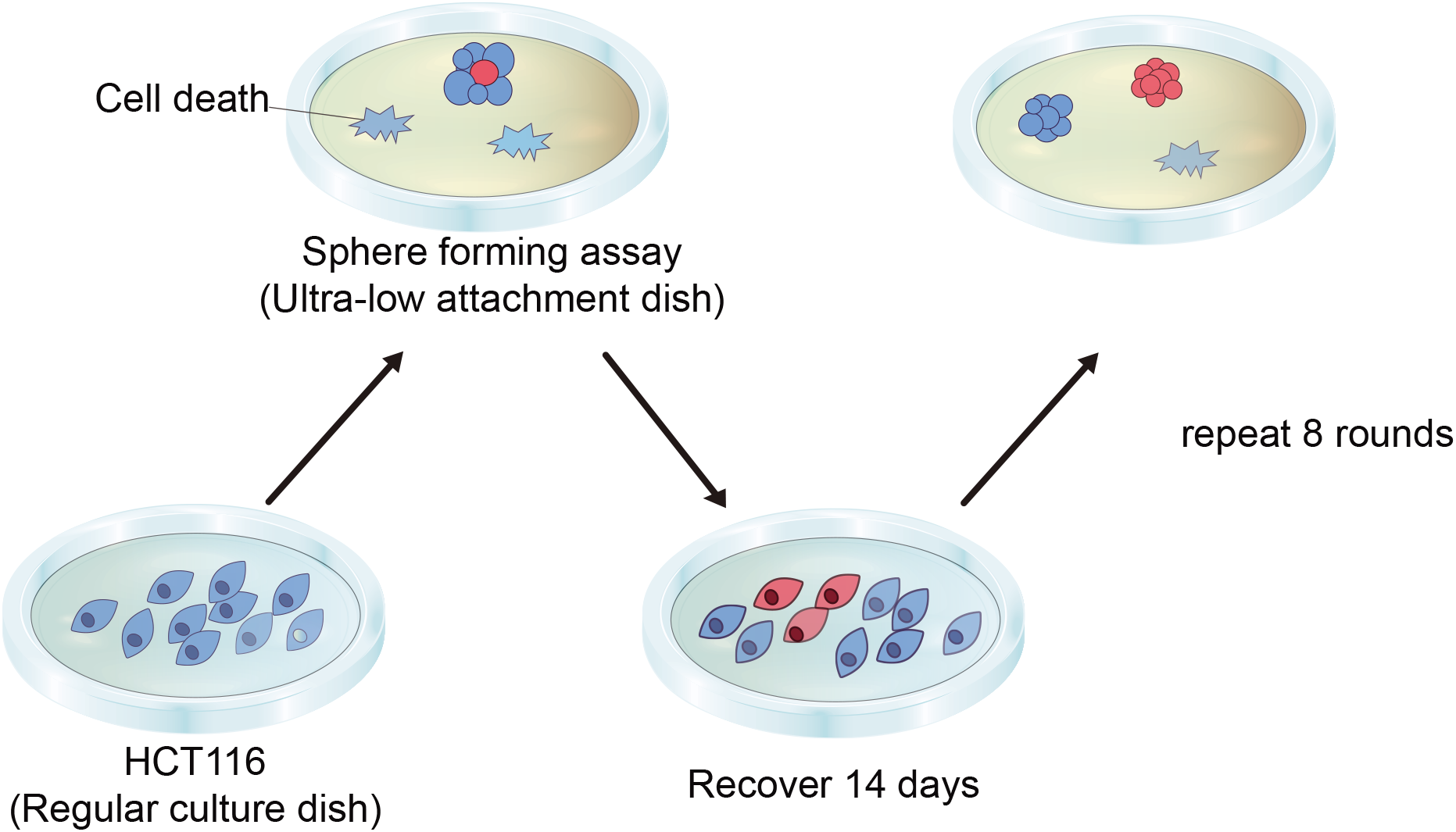
Schematic of the repeated sphere-forming assay for generating sphere-derived cancer stem cell-like cells (SDCSCs) and sphere-derived adherent cells (SDACs) In the sphere-forming assay, the majority of cells die at the beginning of the selection process. Spheres were dissociated to obtain single cells that were re-plated onto a regular dish to recover for 14 days. After recovery, cells were cultured in an ultra-low attachment dish to form spheres. Orange dish: regular culture dish; green dish: ultra-low attachment dish.

HCT116 cells, a human CRC cell line, were used as our model cell line. At the beginning of our assay, HCT116 cells were subjected to a one-time homogenizing sphere-forming assay and re-plated as adherent cells. Such homogenized HCT116 cells were considered parental and designated as ‘G0’. Beginning with this parental line, we performed repeated sphere-forming assay and recovery procedures for the generation of SDCSCs and SDACs. Eighth-generation SDCSCs and SDACs are designated G8S and G8D, respectively.

### Canonical stemness markers are transiently expressed in SDCSCs

To validate that the SDCSCs generated from our sphere-forming assay were CSC-like tumorspheres, we tested the expression of several canonical stemness markers, including the pluripotent markers Nanog and Oct4^18^, and a stem cell marker of the intestinal epithelium, Lgr5^19^. SDACs and SDCSCs from G0 (parental) to G8 were collected. The quantitative real-time polymerase chain reaction (qRT-PCR) revealed that the mRNA expression of these three markers was significantly higher in SDCSCs at every generation than at G0 (Figure 2A). Similar results were observed by immunofluorescent staining of the marker proteins at G1 and G8 (Figure 2B). Up-regulation of canonical markers in SDCSCs validated our repeated sphere-forming assay. The expression of these three markers, however, appeared to be only transient, as their levels were close to that of G0 in SDACs at every generation (Figure 2A and B).

**Figure 2.**
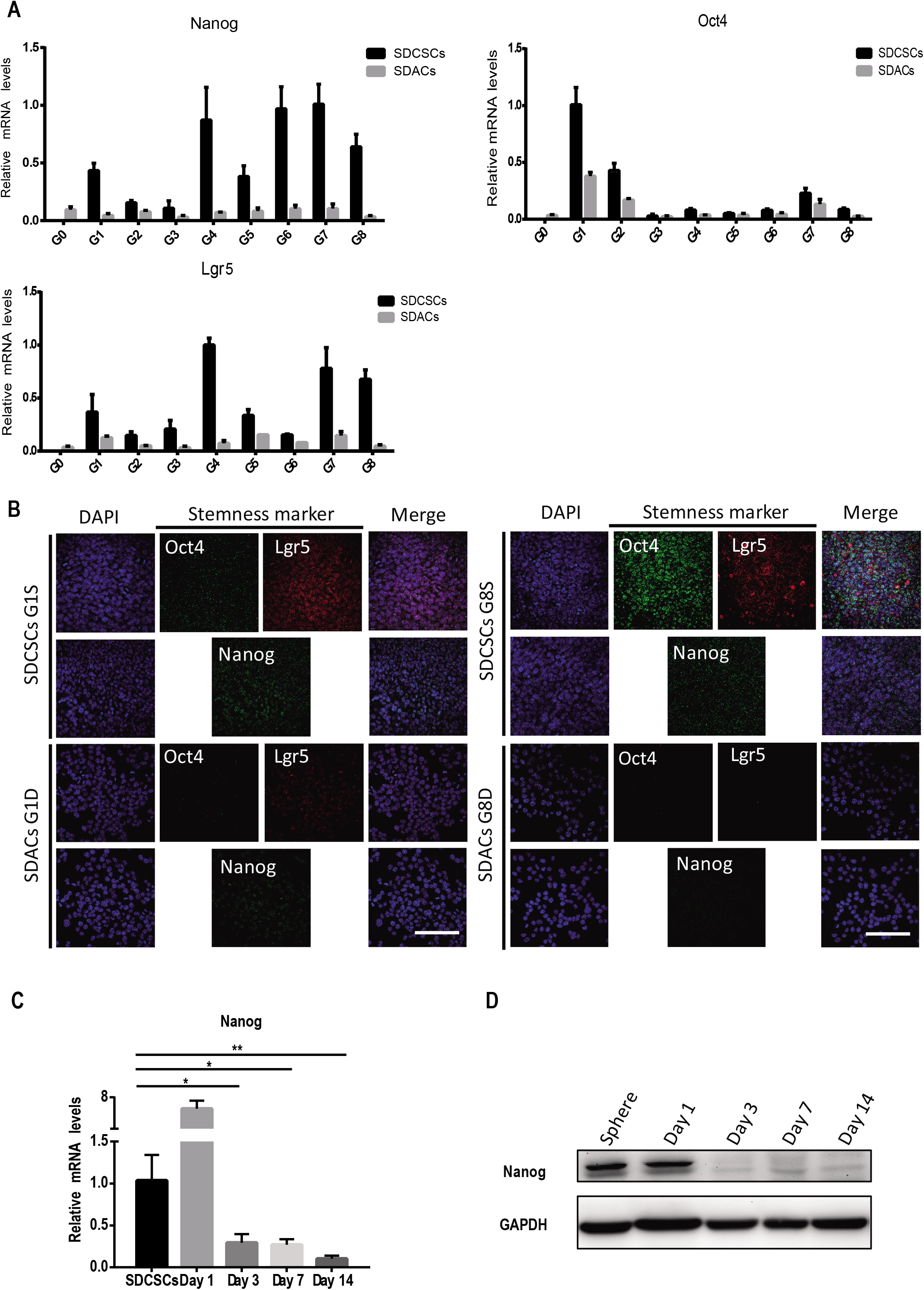
Sphere-derived cancer stem cell-like cells (SDCSCs) express the stemness markers Nanog, Oct4, and Lgr5, transiently. **(A)** Expression of Nanog, Oct4, and Lgr5 mRNAs of SDCSCs and sphere-derived adherent cells (SDACs) from generation G0 (parental) to G8 were analyzed by the quantitative real-time polymerase chain reaction (qRT-PCR). Data are presented as means ± S.D. (n = 3 independent experiments). GAPDH was used as the reference gene. **(B)** Representative immunofluorescent images of SDCSCs (top) and SDACs (bottom) at G1 (left) and G8 (right). Oct4 and Nanog: green; Lgr5: red; DNA: blue; Scale bar: 10 μm. **(C)** RNAs were collected from SDCSCs at days 1, 3, 7, and 14 after re-plating as SDACs. The expression of Nanog mRNA was analyzed by the qRT-PCR. Data are presented means ± S.D. (n = 3 independent experiments). GAPDH was used as the reference gene. Significant differences were determined using ANOVA followed by Dunnett’s test, *p < 0.05, **p < 0.01. **(D)** Proteins were collected from SDCSCs at days 1, 3, 7, and 14 after re-plating as SDACs. Expression of the Nanog protein was detected with western blot. GAPDH was used as the internal control.

Because the expression patterns of Nanog were the most robust among the three markers, we chose to focus on it for investigating the observed transient expression behavior. To analyze the stability of Nanog expression, we performed a time-course experiment. G1S formed spheres, which were collected and re-plated onto a regular dish as adherent cells to be harvested at days 1, 3, 7, and 14. The expression of Nanog was high in SDCSCs and remained high at day 1 in SDACs, but gradually decayed from days 3 to 14 in SDACs, at both the mRNA and protein levels (Figure 2C and D). Such a transient pattern suggests that up-regulation of these markers may stem from the sphere state (i.e., a consequence of the sphere-forming assay) and was not a stable phenotype of the cells.

### Sphere-forming capacity increases significantly after eight rounds of selection

To test whether eight rounds of selection resulted in a cell line that was phenotypically more malignant, we compared the sphere-forming capacities of parental and eighth generation cells *in vitro*^8^. While the morphology of SDACs was similar between G1D and G8D, the size of the spheres was significantly greater in G8S than in G1S (Figure 3A and B, right). The number of spheres that formed was also higher at G8 (Figure 3B, left).

**Figure 3.**
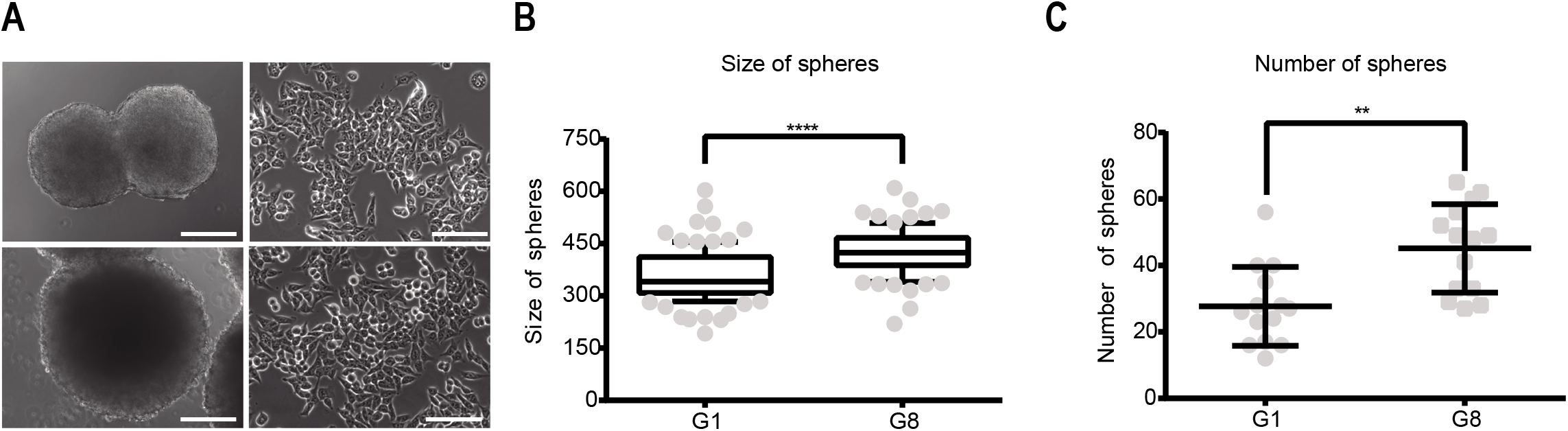
Increased sphere-forming capacity at generation eight (G8) **(A)** Representative phase-contrast images of sphere-derived cancer stem cell-like cells (SDCSCs) (left) and sphere-derived adherent cells (SDACs) (right) at G1 (top) and G8 (bottom). SDCSCs isolated from the sphere-forming assay were allowed to recover for 14 days to obtain SDACs. Scale bar: 100 μm (SDCSCs) and 50 μm (SDACs). **(B and C)** Quantification of the number and size of spheres at G1 and G8. In every condition, 1000 cells were seeded. After 14 days of sphere-formation, the number and size (diameter) of spheres were quantified with ZEN software (n > 120 cells analyzed in both G1 and G8). Data are presented as means ± S.D. Significant differences were determined using Student’s *t* test, **p < 0.01, ****p < 0.0001.

### RNA sequencing identifies DPEP1 as a highly expressed gene in the selected clone

The greater sphere-forming capacity at G8 suggests that our repeated sphere-forming assay was able to select for a more malignant subpopulation (Figure 3A and B). Therefore, we searched for stably up-regulated genes in the selected clone that might be responsible for the malignancy. RNA sequencing was performed to identify genes differentially expressed in G8D as compared to G0 (Figure 4A), and DPEP1was showed to be up-regulated to the greatest extent among the protein-coding genes when the differential expression in the sphere state was included as a reference control (Figure 4B). The up-regulation of DPEP1 mRNA in G8D was confirmed by qRT-PCR (Figure 4C). Because DPEP1 expression is higher in paired tumors than in adjacent tissues^20^ and in tumor tissues than in normal mucosa^21, 22^, we further studied the role of this gene in sphere formation. Two different lentiviral small hairpin RNAs were used to stably knockdown DPEP1 in G8D. Knockdown efficiency was confirmed at both the mRNA and protein levels (Figure 4D and E). Inhibition of DPEP1 led to a marked decrease of the sphere-forming capacity in G8D (Figure 4F and G), suggesting that over-expression of DPEP1 promotes malignancy.

**Figure 4.**
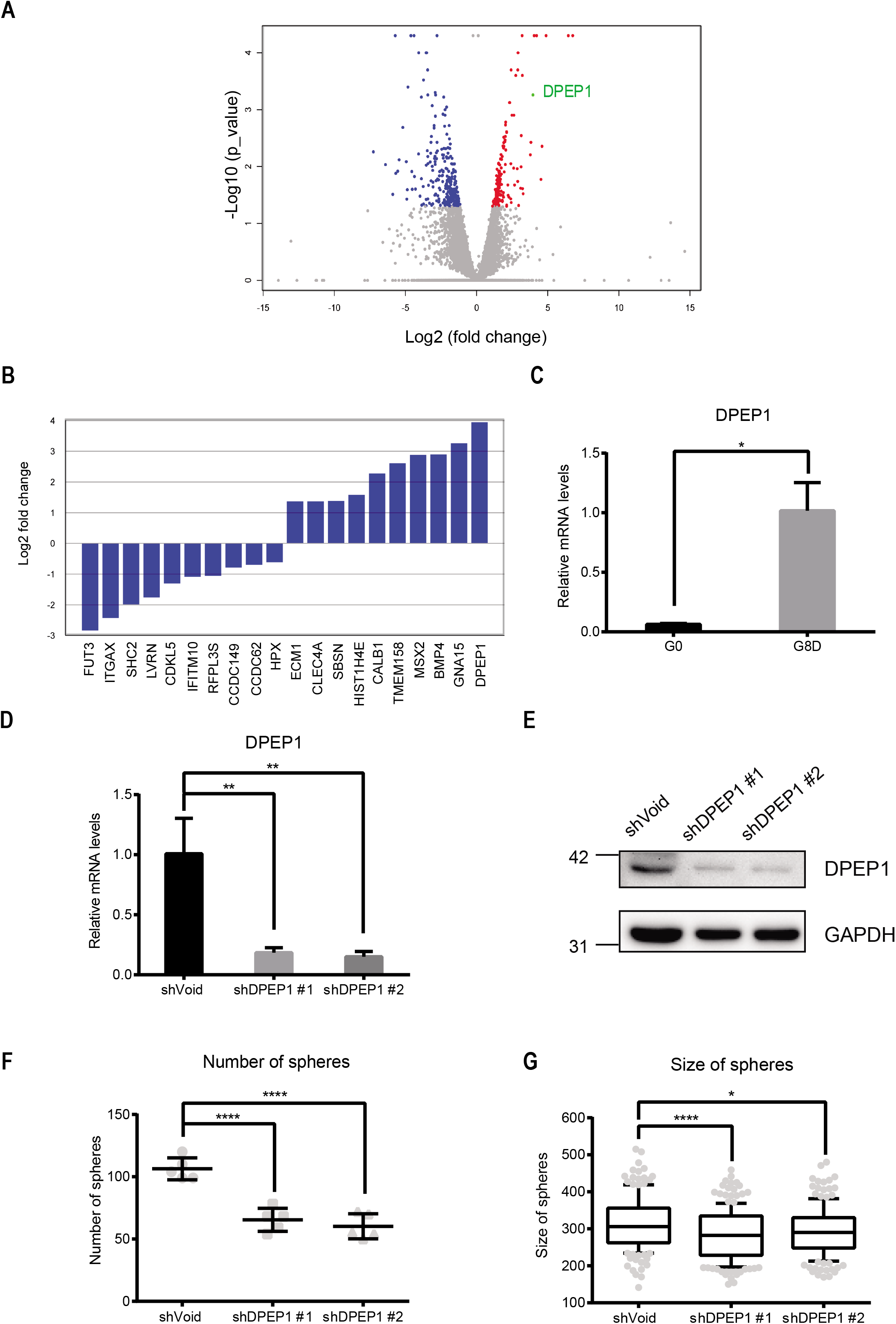
Dipeptidase 1 (DPEP1) promotes sphere-formation in eighth-generation sphere-derived adherent cells (G8D) **(A)** Volcano plot when comparing G8D and G0 with RNA-seq. DPEP1 (marked in green) ranked as one of the significantly upregulated gene among all probes. **(B)** DPEP1 ranked as the top upregulated protein-coding gene. Top 10 up- or down-regulated protein-coding genes were selected based on the criteria: G8S>G1S>G0 AND G8D>G0. In other words, a differential expression in the sphere state was also applied for this selection. **(C)** Differential expression of DPEP1 mRNA between G0 (parental) to G8 was confirmed by the quantitative real-time polymerase chain reaction (qRT-PCR). Data are presented as means ± S.D. (n = 3 independent experiments). GAPDH was used as the reference gene. Significant differences were determined using Student’s *t* test, *p < 0.05. **(D and E)** Knockdown of DPEP1 mRNA **(D)** and protein **(E)** was confirmed by qRT-PCR and western blot, respectively. GAPDH was used as the internal control. Short hairpin (sh)Void: non-targeting negative control shRNA; shDPEP1 #1/#2: two independent shRNAs targeting DPEP1. mRNA data are presented as means ± S.D. (n = 3 independently experiments). Significance differences were determined using ANOVA followed by Dunnett’s test, **p < 0.01. **(F and G)** Quantification of the number and size of spheres following treatment with shVoid or shDPEP1 #1/#2. In every condition, 1000 cells were seeded. After 14 days of sphere-formation, the number and size (diameter) of spheres were quantified with ZEN software (n > 120 cells analyzed in both G1 and G8). Data are presented as means ± S.D. Significant differences were determined using ANOVA followed by Tukey’s test, *p < 0.05, ****p < 0.0001.

### The selected clone exhibits higher DPEP1-dependent tumorgenicity *in vivo*

To investigate whether the selected clone possessed higher DPEP1-dependent tumorgenicity *in vivo*, tumor growth was examined in a xenograft model with severely immunodeficient NSG (NOD/SCID/gamma; i.e., lacking T/B/NK cells) mice^23^. As shown in Figure 5A, tumor size was significantly larger with G8D than with G0 clones, indicating a stronger tumorgenicity after eight rounds of selection. The ablation of DPEP1 greatly reduced tumorgenicity in G8D to that of G0 (Figure 5A; quantification of tumor weights and volumes is shown in Figure 5B and C). Taken together, these data confirm that our repeated sphere-forming strategy can successfully select a highly tumorigenic clone. DPEP1 was stably over-expressed in G8D and responsible for the elevated tumorgenicity.

**Figure 5.**
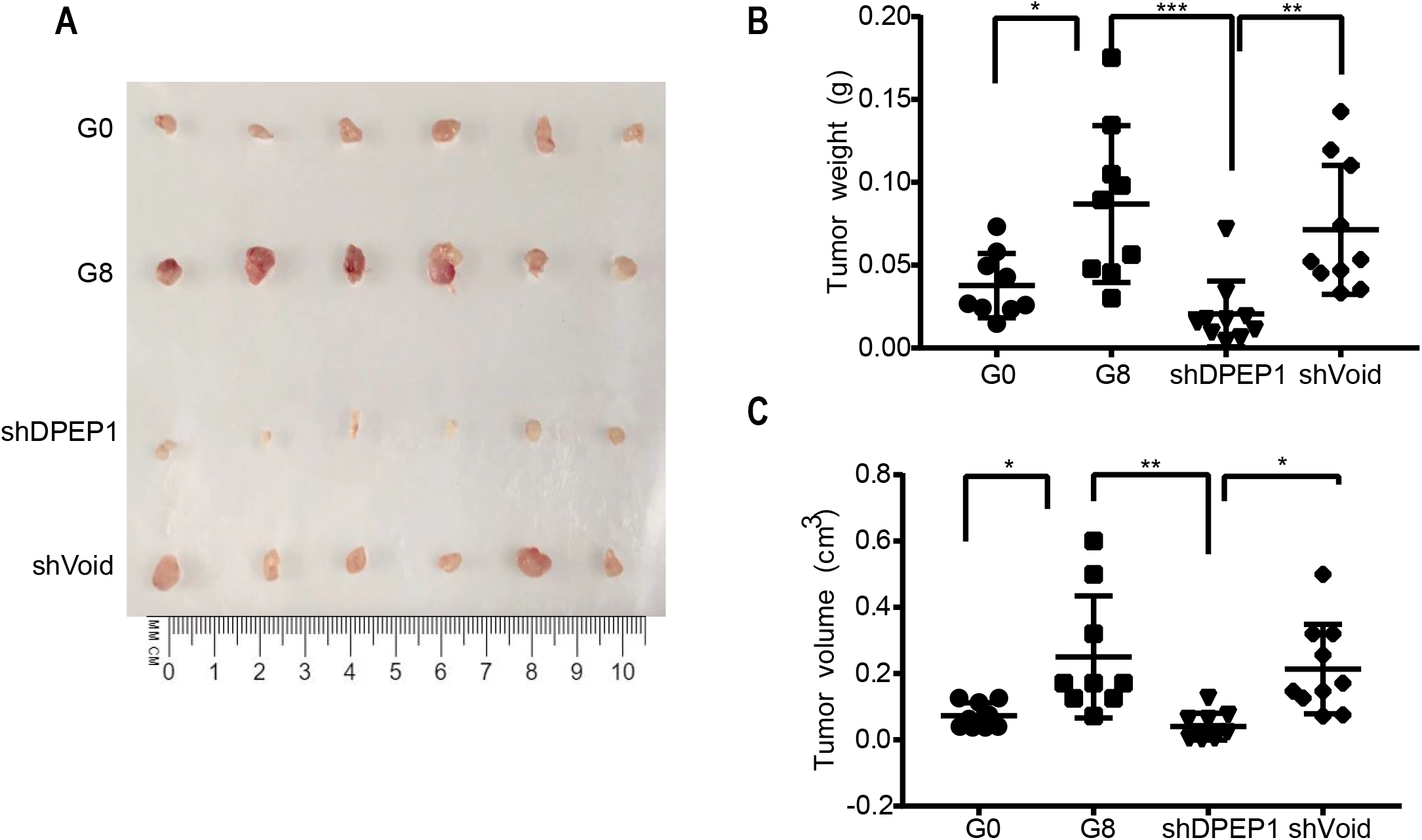
Eighth-generation sphere-derived adherent cells (G8D) exhibit higher dipeptidase 1 (DPEP1)-dependent tumorgenicity *in vivo*. **(A)** Representative images of excised tumors. Cells (1 × 10^2^) were injected subcutaneously into 6-week-old male severely immunodeficient NSG mice. All mice were killed and tumors were excised 28 days after injection. **(B)** Quantification of excised tumor weights for the indicated treatments: G0 (n = 9), G8 (n = 9), G8-short hairpin (sh)DPEP1 (n = 10), and G8-shVoid (non-targeting negative control shRNA) (n = 10). Significant differences were determined using ANOVA followed by Tukey’s test, *p < 0.05, **p < 0.01, ***p < 0.001. **(C)** Tumor volumes were calculated according to the formula: ((*π*/6) × width^2^ × length) for the indicated treatments: G0 (n = 9), G8 (n = 9), G8-shDPEP1 (n = 10), and G8-shVoid (n = 10). Significant differences were determined using ANOVA followed by Tukey’s test, *p < 0.05, **p < 0.01.

## Discussion

Stem cells are defined by their ability to self-renew and differentiate into a variety of cell types. The concept has expanded from embryonic stem cells, to adult stem cells, to CSCs. The sphere-forming assay is used traditionally to identify cells that possess stem cell characteristics, and it has also been adopted to characterize CSCs. Previously, when cancer cells were passaged serially as tumorspheres (i.e., CSC-like cells), later generations were more tumorigenic than parental cells^24^. However, because cells in tumorspheres are cultured temporarily in a condition that is apart from their usual condition, it is likely that the phenotypes measured merely reflect an acute state temporarily. In other words, the expression of markers and phenotypes may be only transient^14^. Indeed, with the three canonical stem cell markers Nanog, Oct4, and Lgr5, we showed that, while they were up-regulated in tumorspheres (SDCSCs), the level decreased to close to the parental level when the spheres were re-plated as adherent cells (SDACs) (Figure 2). Based on this finding, we adopted an alternative “intermittent” strategy. Specifically, after 14 days of sphere formation, cells were allowed to rest in the adherent state for 14 days before the next sphere-forming assay (Figure 1). With this experimental strategy, we generated cell lines and identified stable phenotypes. Another benefit of the strategy was that cells could be expanded and stored in the resting SDAC state, and could be recovered at every generations. As opposed to continuous propagation as tumorspheres^24^, which only lasts for a limited duration in our hands (data not shown), in theory, with the intermittent strategy, the assay can be carried out for infinite generations.

With this intermittent strategy, we were able to isolate a clone after eight rounds of sphere formation that, in its adherent state, was more tumorigenic than the parental line (Figure 5). Because the number of selection rounds was relatively small, it is probable that the selected clone (G8D) was a preexisting subpopulation within the heterogeneous parental line. However, given the genomic instability of cancer cells, it is likely that increasing the number of selections would allow accumulation of new mutations during the course of the experiment. Thus, the system has the potential to be used to carry out “experimental cancer evolution”^25^. In such a case, the hypoxic microenvironment within the tumorsphere can serve as the selection pressure that drives evolution towards a potentially more devastating phenotype^26^. Given the previous success in experimental evolution of multicellularity (i.e., stemness) from unicellular *Saccharomyces cerevisiae*^27^ and *Chlamydomonas*^28^, it seems reasonable that we may be able to experimentally evolve stable stemness in cancer. Such a stable CSC-like line would be a good experimental model for drug screening for anti-CSC agents.

In our transcriptome analysis of the selected clone, we identified DPEP1 as the main differentially expressed gene (Figure 4A). We also showed that DPEP1 was responsible for the elevated sphere-forming capacity *in vitro* (Figure 4E and F) and tumorigenicity *in vivo* (Figure 5A-C). These findings coincide with at least three previous studies that reported DPEP1 was highly expressed in malignant colorectal tissues^20–22^. Because a high DPEP1 level is associated with poor CRC patient survival^21^, our results further strengthen the concept of using DPEP1 expression as a CRC prognostic marker. Mechanistically, it has been reported that DPEP1 promotes cell proliferation *in vitro*^20^. Because DPEP1 is a matrix metalloproteinase, it may also promote tumor growth *in vivo* by degrading matrix barriers to enhance cell migration and angiogenesis^29^. Further investigations are necessary to disentangle the roles of DPEP1 that may contribute to *in vivo* tumor growth.

## Material and Methods

### Cell culture, sphere-forming assay, and preparation of SDACs

The human CRC line, HCT116, was obtained from the Biosource Collection and Research Center (Taiwan) and maintained in McCoy’s 5A medium (Gibco) supplemented with 10% fetal bovine serum (HyClone) and 1% penicillin/streptomycin (Gibco) at 37°C in a humidified 5% CO2 incubator. For sphere formation, cells were plated at 1000 cells/well in 24-well ultra-low attachment plates (Corning) and cultured in stem cell medium for 14 days. This medium consisted of serum-free Dulbecco’s modified Eagle’s medium (DMEM)/F12 (1:1; Gibco) supplemented with B27 (Gibco), minimum essential medium non-essential amino acids, sodium pyruvate, L-glutamine, insulin (Gibco), 20 ng/mL epidermal growth factor, 10 ng/mL basic fibroblast growth factor, and 100 U/mL penicillin/streptomycin.

To prepare SDACs, TrypLE Express (Gibco) was used to dissociate spheres, and single cells were collected by centrifugation. The cells were then resuspended in DMEM/F12 medium supplemented with 10% fetal bovine serum and 1% penicillin/streptomycin for 14 days before the next sphere-forming assay round.

### Microscopy and immunofluorescence staining

Phase-contrast images of SDCSCs and SDACs were obtained using an inverted Axio Observer Z1 microscope (10×/0.30, 63×/0.75) with a CoolSnap HQ2 camera and ZEN imaging software (Carl Zeiss). For immunofluorescence and confocal imaging, cells were fixed with 3.7% formaldehyde (Sigma-Aldrich) for 20 min at 37°C, permeabilized with 0.3% Triton X-100 (Sigma-Aldrich) in PBS for 5 or 20 min at room temperature, blocked in 1% bovine serum albumin (BSA) (Sigma-Aldrich) in PBS for 30 min at room temperature, and incubated in 1% BSA/PBS with fluorescein isothiocyanate-conjugated anti-α-tubulin (1:200) (Sigma-Aldrich), Lgr5 (1:200) (Origene, TA503316), Oct4 (1:200) (Santa Cruz Biotechnology, SC-9081), and Nanog (1:50) (Santa Cruz, SC-293121) antibodies for 1 h at room temperature. Slides were washed three times in PBS, then incubated in 1% BSA/PBS with 1:500 goat anti-mouse (Thermo Fisher, A21424) or goat anti-rabbit (Abcam, ab150077) IgG H&L. ProLong Gold Antifade Mountant with 4′,6-diamidino-2-phenylindole (Life Technologies) was used for mounting. Images were obtained with a Leica TCS SP5 confocal microscope using a HCX PL APO lambda blue 63.0 × 1.40 oil immersion objective lens (Applied Precision). Fluorescence excitation, selected with an acousto-optic tunable filter, was at 405 nm from a diode laser for 4′,6-diamidino-2-phenylindole, 488 nm from an argon laser for Oct4, and 561 nm from a DPSS laser for Nanog and Lgr5. Emission was measured at 500-550 nm. Image acquisition was controlled using LAS AF Lite software (https://microscopy.duke.edu/image-analysis). Acquisition of multiple stage positions used a Z-Galvo stage.

### RNA extraction and qRT-PCR

Total RNA was extracted with phenol (Sigma-Aldrich)/chloroform (Avantor) and converted to cDNA using the iScript cDNA Synthesis Kit (BioRad). QRT-PCR reactions were carried out with a SYBR Green kit (Kapa Biosystems) on a CFX96 qRT-PCR machine (BioRad). Relative mRNA levels were calculated according to the ΔΔCt method. GAPDH was used as the housekeeping gene. The primer sequences were:

GAPDH forward: 5′-TGG TGA AGC AGG CGT CGG AG-3′; reverse: 5′-GGT GGG GGA CTG AGT GTG GC-3′
*β*-Actin forward: 5′-AGA GCT ACG AGC TGC CTG AC-3′; reverse: 5′-AGC ACT GTG TTG GCG TAC AG-3′
OCT4 forward: 5′-AGC TTG GGC TCG AGA AGG AT-3′; reverse: 5′-AGA GTG GTG ACG GAG ACA GG-3′
NANOG forward: 5′-ACA ACT GGC CGA AGA ATA GCA-3′; reverse: 5′-GGT TCC CAG TCG GGT TCA C-3′
LGR5 forward: 5′-CTC CCA GGT CTG GTG TGT TG-3′; reverse: 5′-GAG GTC TAG GTA GGA GGT GAA G-3′
DPEP1 forward: 5′-CAA GTG GCC GAC CAT CTG G-3′; reverse: 5′-GGG ACC CTT GGA ACA CCA TC-3′

### RNA sequencing

RNA concentrations and purity were determined by measuring the OD260/OD280 (>1.8) and OD260/OD230 (>1.6), respectively. Yield and quality were assessed using an Agilent 2100 Bioanalyzer (Agilent Technologies). After the sample quality control procedures, mRNA from eukaryotic organisms was enriched using oligo(dT) beads. First, mRNA was fragmented randomly by adding fragmentation buffer, then cDNA was synthesized using an mRNA template and random hexamer primers, after which a custom second-strand synthesis buffer (Illumina), dNTPs, RNase H, and DNA polymerase I were added to initiate second-strand synthesis. Second, after a series of terminal repair reactions, ligation, and sequencing adaptor ligation, the double-stranded cDNA library was completed through size selection and PCR enrichment. The libraries were pooled and analyzed on an Illumina sequencer using the paired-end 150 bp RapidRun format to generate 20 million total reads per sample. Raw reads of RNA-seq from the sequencing instrument first trimmed the low-quality tranche and were checked. Spliced Transcripts Alignment to a Reference software (Illumina) was used to map spliced short-read (RNA-seq reads) to the reference genome (Ensembl GRCh38). Based on spliced alignments, transcript reconstruction and estimations of transcript abundance were conducted by Cuffquant. Gene expression was normalized by calculating the number of RNA-seq fragments per kilobase of transcript per total million fragments mapped. Cuffdiff was used to test the statistical significance of observed changes and identify genes that were differentially regulated at the transcriptional or post-transcriptional levels.

### Western blot

Cells were washed with ice-cold PBS. Total cell lysates were extracted by radioimmunoprecipitation assay buffer (Millipore) supplemented with protease and phosphatase inhibitors (Roche). Protein concentrations were determined using a Bradford protein assay kit (BioRad). Exactly 30 or 100 μg of protein were separated on 10% polyacrylamide gels and then transferred onto membranes. The membranes were washed in PBS with Tween 20 (PBST). After blocking with 5% nonfat milk in PBST for 1 h at room temperature, the membranes were incubated with anti-Nanog (Santa Cruz Biotechnology, SC-293121), anti-DPEP1 (Signalway Antibody, #38797), and anti-GAPDH (GeneTex, GTX627408) antibodies overnight at 4°C. After incubation with the corresponding secondary antibody for 1 h at room temperature, immunoreactive proteins were detected by an enhanced chemiluminescence detection system (EMD-Millipore).

### Xenograft tumorigenicity assay

Xenograft tumorigenicity was determined as described previously. Briefly, HCT116 cells at different tumorsphere generations (G0 or G8D) or treatments (G8D with shVoid (non-targeting negative control shRNA) or shDPEP1 viral infection) were harvested, washed with PBS, and resuspended in DMEM/F12 medium. Cells (1 × 10^2^) were then injected subcutaneously into the right and left flank regions of 6-week-old male NSG mice (Genomic Research Center, Taiwan). All mice were killed 28 days after injection, and tumors were surgically excised, weighed, the volume measured, and photographed. Differences in tumor progression were analyzed statistically by an analysis of variance followed by Dunnett’s or Tukey’s *post-hoc* test. P < 0.05 was considered significant.

### Author contributions

P.-L. L. performed the animal experiments and analyzed the RNA-seq data. C.-T. C. performed the repeated sphere-forming experiments. C.-Y. F. characterized DPEP1. H. L. performed the animal experiments. Y. C. performed the lentiviral knockdown experiments. Ng. W. and H. M. assisted in the animal experiments. P.-L. L., J. L., and H.-C. H. directed the project, analyzed data, discussed data, and wrote the manuscript.

## Acknowledgements

We thank Technology Commons (College of Life Science, National Taiwan University) for use of the Leica TCS SP5 confocal microscope. We thank the Pathology Core Laboratory (Academia Sinica) for assisting with the animal experiments. This work was funded by the Ministry of Science and Technology (102-2311-B-002-041-MY3) to H.-C. H. and Academia Sinica (AS-SUMMIT-108) to J.L.

## Reference

1. Jemal A, Bray F, Center MM, Ferlay J, Ward E, Forman D. Global cancer statistics. CA Cancer J Clin. 2011;61:69–90.

2. Bhandari A, Woodhouse M, Gupta S. Colorectal cancer is a leading cause of cancer incidence and mortality among adults younger than 50 years in the USA: a SEER-based analysis with comparison to other young-onset cancers. J Investig Med. 2017;65:311–315.

3. Harrison S, Benziger H. The molecular biology of colorectal carcinoma and its implications: a review. Surgeon. 2011;9:200–210.

4. Siegel R, Ward E, Brawley O, Jemal A. Cancer statistics, 2011: the impact of eliminating socioeconomic and racial disparities on premature cancer deaths. CA Cancer J Clin. 2011;61:212–236.

5. Vaiopoulos AG, Kostakis ID, Koutsilieris M, Papavassiliou AG. Colorectal cancer stem cells. Stem Cells. 2012;30:363–371.

6. Dean M, Fojo T, Bates S. Tumour stem cells and drug resistance. Nat Rev Cancer. 2005;5:275–284.

7. Altman J. Autoradiographic and histological studies of postnatal neurogenesis. IV. Cell proliferation and migration in the anterior forebrain, with special reference to persisting neurogenesis in the olfactory bulb. J Comp Neurol. 1969;137:433–457.

8. Pastrana E, Silva-Vargas V, Doetsch F. Eyes wide open: a critical review of sphere-formation as an assay for stem cells. Cell Stem Cell. 2011;8:486–498.

9. Singh SK, Clarke ID, Terasaki M, et al. Identification of a cancer stem cell in human brain tumors. Cancer Res. 2003;63:5821–5828.

10. Ponti D, Costa A, Zaffaroni N, et al. Isolation and in vitro propagation of tumorigenic breast cancer cells with stem/progenitor cell properties. Cancer Res. 2005;65:5506–5511.

11. Ricci-Vitiani L, Lombardi DG, Pilozzi E, et al. Identification and expansion of human colon-cancer-initiating cells. Nature. 2007;445:111–115.

12. Todaro M, Alea MP, Di Stefano AB, et al. Colon cancer stem cells dictate tumor growth and resist cell death by production of interleukin-4. Cell Stem Cell. 2007;1:389–402.

13. Amsterdam A, Raanan C, Schreiber L, et al. Differential localization of LGR5 and Nanog in clusters of colon cancer stem cells. Acta Histochem. 2013;115:320–329.

14. Yakisich JS, Azad N, Kaushik V, Iyer AKV. Cancer Cell Plasticity: Rapid Reversal of Chemosensitivity and Expression of Stemness Markers in Lung and Breast Cancer Tumorspheres. J Cell Physiol. 2017;232:2280–2286.

15. Chu YW, Yang PC, Yang SC, et al. Selection of invasive and metastatic subpopulations from a human lung adenocarcinoma cell line. Am J Respir Cell Mol Biol. 1997; 17:353–360.

16. Yang CF, Tsai WY, Chen WA, et al. Kinesin-5 Contributes to Spindle-length Scaling in the Evolution of Cancer toward Metastasis. Sci Rep. 2016;6:35767.

17. Chen JJ, Peck K, Hong TM, et al. Global analysis of gene expression in invasion by a lung cancer model. Cancer Res. 2001;61:5223–5230.

18. Liu A, Yu X, Liu S. Pluripotency transcription factors and cancer stem cells: small genes make a big difference. Chin J Cancer. 2013;32:483–487.

19. de Sousa e Melo F, Kurtova AV, Harnoss JM, et al. A distinct role for Lgr5(+) stem cells in primary and metastatic colon cancer. Nature. 2017;543:676–680.

20. Hao JJ, Zhi X, Wang Y, et al. Comprehensive Proteomic Characterization of the Human Colorectal Carcinoma Reveals Signature Proteins and Perturbed Pathways. Sci Rep. 2017;7:42436.

21. Eisenach PA, Soeth E, Roder C, et al. Dipeptidase 1 (DPEP1) is a marker for the transition from low-grade to high-grade intraepithelial neoplasia and an adverse prognostic factor in colorectal cancer. Br J Cancer. 2013;109:694–703.

22. Park SY, Lee SJ, Cho HJ, et al. Dehydropeptidase 1 promotes metastasis through regulation of E-cadherin expression in colon cancer. Oncotarget. 2016;7:9501–9512.

23. Shultz LD, Lyons BL, Burzenski LM, et al. Human lymphoid and myeloid cell development in NOD/LtSz-scid IL2R gamma null mice engrafted with mobilized human hemopoietic stem cells. J Immunol. 2005;174:6477–6489.

24. Cao L, Zhou Y, Zhai B, et al. Sphere-forming cell subpopulations with cancer stem cell properties in human hepatoma cell lines. BMC Gastroenterol. 2011;11:71.

25. Taylor TB, Johnson LJ, Jackson RW, Brockhurst MA, Dash PR. First steps in experimental cancer evolution. Evol Appl. 2013;6:535–548.

26. Yun Z, Lin Q. Hypoxia and regulation of cancer cell stemness. Adv Exp Med Biol. 2014;772:41–53.

27. Ratcliff WC, Denison RF, Borrello M, Travisano M. Experimental evolution of multicellularity. Proc Natl Acad Sci U S A. 2012;109:1595–1600.

28. Ratcliff WC, Herron MD, Howell K, Pentz JT, Rosenzweig F, Travisano M. Experimental evolution of an alternating uni- and multicellular life cycle in Chlamydomonas reinhardtii. Nat Commun. 2013;4:2742.

29. Deryugina EI, Quigley JP. Matrix metalloproteinases and tumor metastasis. Cancer Metastasis Rev. 2006;25:9–34.

